# Opposing effects of slow and fast theta synchrony on working memory in the human hippocampal-orbitofrontal network

**DOI:** 10.64898/2026.05.10.724153

**Authors:** Samantha M. Gray, Adam J. O. Dede, Zachariah R. Cross, Ignacio Saez, Fady Girgis, Edward F. Chang, Kurtis I. Auguste, Ammar Shaikhouni, Robert T. Knight, Elizabeth L. Johnson

**Affiliations:** Northwestern University, Department of Medical Social Sciences; Stanford University, Department of Psychiatry and Behavioral Sciences; La Trobe University, Department of Psychology and Public Health; Icahn School of Medicine at Mount Sinai, Departments of Neuroscience, Neurosurgery, and Neurology; University of California, Davis, Department of Neurology and Center for Mind and Brain; University of Calgary, Department of Clinical Neurosciences; University of California, San Francisco, Department of Neurological Surgery, and UCSF Benioff Children’s Hospital, Department of Pediatric Surgery; Ohio State University, Department of Neurological Surgery, and Nationwide Children’s Hospital; University of California, Berkeley, Helen Wills Neuroscience Institute and Departments of Neuroscience and Psychology; Northwestern University, Departments of Psychology and Pediatrics

## Abstract

Working memory (WM) enables us to maintain and manipulate information over time, but how the brain organizes sequential information locally and across networks remains unclear. Recent work suggests that slow and fast theta oscillations serve different roles in memory, yet their distinct contributions to sequential WM are unknown. Based on evidence that the hippocampus (HC) and orbitofrontal cortex (OFC) support sequential WM and that slower theta cycles provide optimal temporal windows for organizing items in WM, we predicted that these regions would coordinate via slow theta dynamics. We analyzed intracranial EEG from the HC, OFC, and amygdala (AMY) in 21 neurosurgical patients (7 female, 13-54 years of age; M ± SD, 30 ± 11.2 years) performing a delayed match-to-sample WM task. We assessed phase locking between regions, phase-amplitude coupling within regions, and neuronal phase coding for slow (~1-4.5 Hz) and fast (~4.5-8 Hz) theta oscillations. We found significant slow and fast theta synchrony between all regions, but identical anatomical pathways produced opposing behavioral effects depending on oscillatory frequency, particularly during higher cognitive demand. Slow theta synchrony was associated with faster response times (RTs), while fast theta synchrony between HC and OFC hindered both accuracy and RTs. Unexpectedly, AMY modulated RT through demand-dependent slow theta synchrony, where AMY-OFC synchrony predicted faster RTs during maintenance and HC-AMY synchrony predicted faster RTs during higher cognitive demand. Sustained coupling between slow theta oscillations and high-frequency broadband activity within each region suggests that local organization coincides with beneficial network behavioral effects. These results establish a frequency-opponent mechanism in which theta oscillation frequencies determine whether HC-OFC circuits facilitate or impair sequential WM.

## Introduction

Working memory (WM) is a crucial component of cognition that enables the maintenance and manipulation of spatial, temporal, and semantic information in mind. The ramifications of disrupted WM are evident in traumatic brain injury and virtually all neuropsychiatric disorders^1–5^, underscoring its importance to daily life. A key open question in WM is how we parse and maintain incoming streams of information while accessing individual objects and their features within a stream^6^. Studies of WM using the Sternberg task have demonstrated that response time (RT) scales with the number of items held in memory, suggesting that mnemonic representations are accessed serially^7^. Yet, WM must rely on precisely binding a given stimulus to its spatial location and temporal position relative to other stimuli given our ability to maintain multiple features of a given stimulus^8–10^. This binding requirement presents significant computational challenges, requiring the brain to track multiple dimensions of information while maintaining their relationships across time and integrating new information. The neural mechanisms underlying the process of actively maintaining sequential information and subsequently integrating new information to execute behavioral judgements remain undefined. Developing a mechanistic understanding of how humans encode, maintain, and compare sequential information in mind across distributed neural circuits could facilitate the development of neurotherapeutic interventions for WM deficits in neuropsychiatric conditions.

Activity in frontal and medial temporal brain regions such as the hippocampus (HC) have been shown to facilitate WM processes. Early work in non-human primates demonstrated persistent firing of frontal neurons during stimulus maintenance in the absence of external stimuli as a hallmark of WM^11,12^. In humans, studies using stereotactic EEG (sEEG) further demonstrated persistent firing of HC neurons during WM maintenance in the absence of external stimuli13, implicating both frontal and HC regions in WM. Still, the extent to which HC contributes to WM remains debated, with conflicting evidence from lesion and neuroimaging studies^13^. Greater activity in the amygdala (AMY), a region traditionally associated with emotion processing, has also been linked to faster RTs in fMRI studies of WM for emotionally neutral information^14^. AMY involvement in non-emotional cognitive tasks is likely due to dense structural and functional connectivity between it and the HC^15^, as well as structural connectivity with orbital regions of the frontal lobe^16^. Indeed, we have reported that the human orbitofrontal cortex (OFC) is causally implicated in WM for sequential information, supported by sequential WM deficits in lesioned patients that were commensurate with OFC lesion size^17^. OFC is structurally connected to the medial temporal lobe via the uncinate fasciculus^18,19^, providing the infrastructure for OFC to rapidly interact with HC and AMY during sequential WM.

In addition to persistent neuronal firing, WM processes have been linked to theta oscillations (~2–8 Hz) across multiple species^20–22^. A dominant model of WM states that individual memory items are represented by discrete gamma cycles (>30 Hz) that are temporally organized within the phase of ongoing theta oscillations^23,24^. Consistent with this model, the modulation of gamma power by theta phase, termed theta-gamma phase-amplitude coupling (PAC), has been shown to support the maintenance of temporal order in WM^23,25–27^. In tasks requiring sequential WM, PAC strength scales with the number of items in a sequence and predicts behavioral performance as indexed by faster RTs^25,26^. Though classically described as theta-gamma coupling, high-frequency broadband activity (HFB; ~70-150 Hz) tracks population-level neuronal firing and hemodynamic responses^28–33^, and has been shown to be functionally relevant in human WM processes^34^. Research has demonstrated theta-HFB PAC in cortical and temporal regions^25,35^, and causal non-invasive work has demonstrated that enhancing theta-HFB PAC in the frontal cortex enhances WM performance^36^. Similar to PAC, a growing body of cross-species research demonstrates that the timing of neuronal spikes relative to the phase of ongoing theta oscillations becomes more structured during sequence encoding^37,38^. This phenomenon, known as phase coding, demonstrates that neurons convey information not only through their firing rate but also the timing of their firing within an oscillatory cycle. Neurons that fire at different phases of the same rhythm have been shown to encode distinct elements of a sequence, spatial locations, and object identities, providing a temporal code for organizing information in WM^37,39–41^. Together, PAC and phase coding provide complementary mechanisms for organizing sequential information within theta cycles, where different theta phases maintain high-frequency neural events in sequences.

Whereas PAC and phase coding organize information within individual brain regions, oscillatory synchrony enables communication and information integration across regions, thus facilitating the coordination necessary for complex cognitive functions such as WM^42–44^. Recent work has suggested that theta synchrony between hippocampal and frontal regions supports sequential WM^45,46^. Previous research has further shown elevated AMY-HC connectivity during the processing of salient stimuli, suggesting that AMY contributes to behaviorally relevant cognitive processing through oscillatory interactions with HC. These findings led us to hypothesize that while nesting high-frequency activity within theta rhythms supports the serial organization of sequential WM items, interregional theta synchrony supports the transfer of sequential representations between frontal and temporal regions. We further hypothesized that frontotemporal coordination supporting sequential WM would be mediated by theta oscillations that differ in frequency, with slow and fast theta serving distinct functional roles.

Recent work has provided evidence for two distinct theta oscillations in frontal and medial temporal regions: a slower oscillation ~1.5-4.5 Hz and faster oscillation ~4.5-8.0 Hz^47–51^. Slow and fast theta frequencies exhibit distinct peaks in the power spectra^47,52^, have different developmental trajectories, and may serve different functional roles in memory processes^47–49,52^. Emerging evidence implicates slow theta in sequential WM processes. Specifically, computational models propose that the longer timescales of slow theta oscillations are better suited for organizing extended temporal sequences and packaging sequential information^23,24^. Recent empirical evidence provides initial support for this proposal by demonstrating PAC between slow, but not fast, theta oscillations and HFB in HC during the maintenance of sequential information^25^. Moreover, studies using transcranial alternating current stimulation (tACS) to induce slow and fast theta oscillations have shown a beneficial role of slow, but not fast, theta enhancement on temporal memory^53–55^. Although HC theta phase coding has been shown to represent sequential order tied to spatial or semantic features (i.e., concept and place cells)^26,28–30^, it is unknown whether similar coding schemes extend to more abstract representations of temporal order in WM, are specific to slow or fast theta, or occur in across the HC-OFC network. We tested the specific hypothesis that sequential WM would be supported by slow theta synchrony between HC and OFC, as well as slow theta-HFB PAC and phase coding within HC and OFC, which together may organize and transfer sequential representations across the circuit.

To test this pathway, we recorded single-neurons and local field potentials (LFPs) from HC, AMY, and OFC in 21 neurosurgical patients as they performed a delayed match-to-sample sequential WM task. On each trial, participants memorized a three-item sequence of colored shapes varying in spatial location on the screen (“sample sequence”), followed by a 2-second delay and a new sequence of three shapes (“test sequence”) (Figure 1A)^46,57,58^. Stimuli were randomized to prevent overlearning, using a pool of 16 shapes and 11 colors (i.e., 176 unique stimuli). This task engages multiple WM subdomains and varies in cognitive demand within a trial, offering a semi-naturalistic framework to probe WM. We examined how the HC, AMY, and OFC coordinate via oscillatory theta mechanisms to support the encoding, maintenance, and comparison of sequential information. We observed slow and fast theta synchrony between all regions, but identical anatomical pathways produced opposing behavioral effects depending on oscillatory frequency, particularly during higher cognitive demand. Slow theta synchrony was associated with faster RTs across the circuit while fast theta synchrony between HC and OFC hindered both accuracy and RTs. Additionally, we reveal sustained slow theta PAC in both HC and OFC throughout the task, suggesting that local organization of sequential information coincides with beneficial network effects. Finally, AMY modulated RT via slow theta synchrony with OFC during lower cognitive demand and HC during higher cognitive demand. Together, these results establish a frequency-opponent mechanism in which theta oscillation frequency determines whether HC-OFC circuits facilitate or impair sequential WM, and point to a novel role for AMY in demand-dependent modulation of response speed.

**Figure 1.**
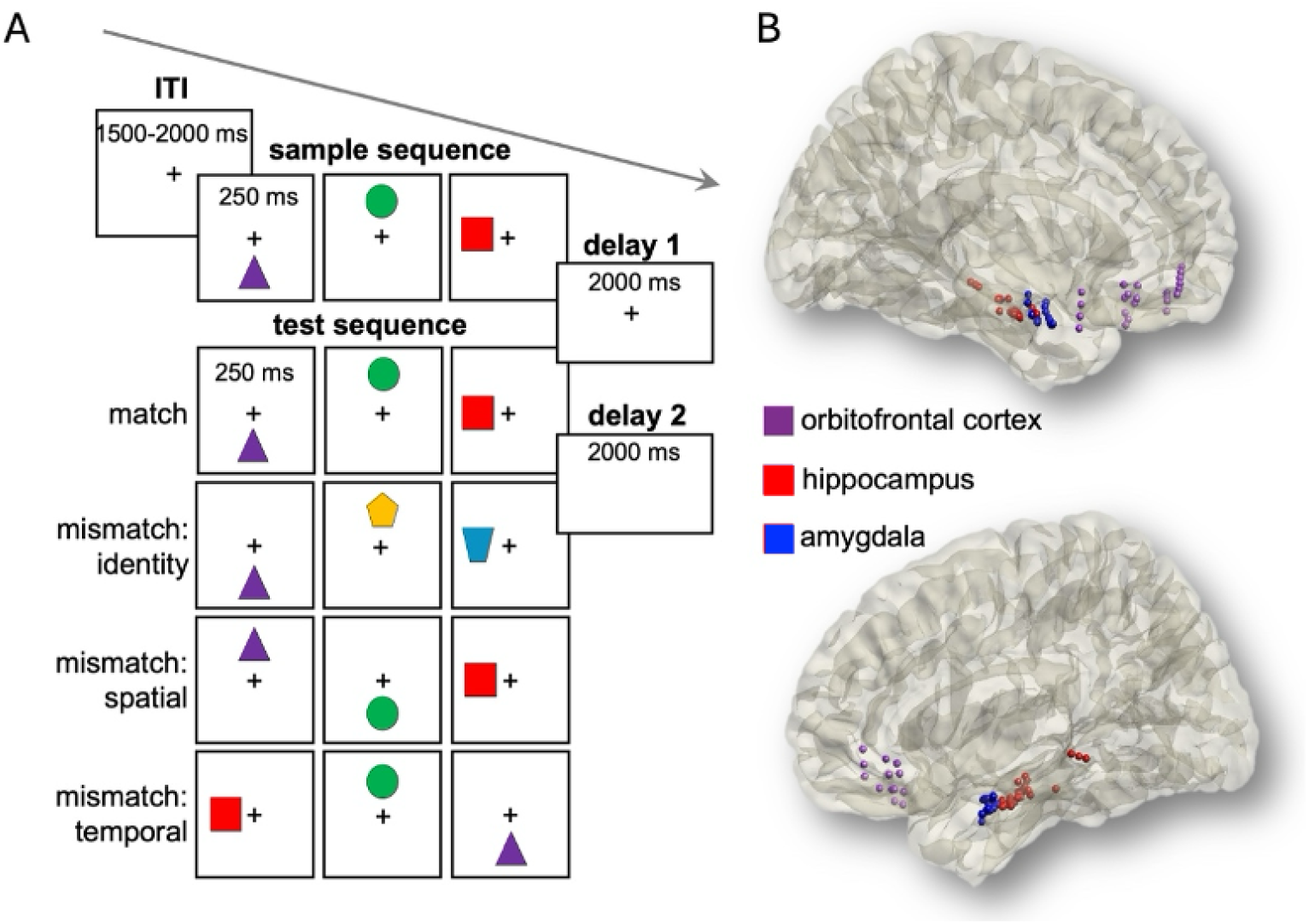
Design and electrode coverage. (A) Subjects completed a WM-DMS task, where they were presented with two sequences of three stimuli (sample and test) separated by a delay and asked to determine if the sample sequence was a match or a mismatch to the test sequence. (B) Artifact- and seizure-free sEEG coverage in OFC (n=36), HC (n=51), and AMY (n=43) across all subjects shown on the MNI-152 template brain.

## Results

### Patients exhibit intact sequential WM performance

Twenty-one neurosurgical patients (7 females) with electrodes implanted in HC (n=51), AMY (n=43), and OFC (n=36) performed the WM-DMS task (see Figure 1A). RT for each patient was recorded per trial (M ± SD: 2383.71 ± 988.91ms) and was measured as the time from the presentation of the match/mismatch question to the point at which participants clicked their response. All patients included in this study had AR of at least 0.73 (see Methods for AR calculation), with performance comparable to healthy controls reported in other studies using the same WM-DMS task (M ± SD: 0.92 ± 0.07, healthy controls M ± SD: 0.85 ± 0.12)^46,57^.

### Theta activity during sequential WM in HC, AMY, and OFC

We first confirmed that theta activity was present during the task by computing time-frequency representations of power from 1 to 200 Hz per ROI (i.e., −0.75 to 6.5 s from sample onset, averaged over trials, channels, and subjects within each ROI). Power was z-scored using statistical bootstrapping^34,44,59–61^. We observed dissociable timing in theta power between the medial temporal regions and OFC (Figure 2), with OFC power at its maximum during sample sequence presentation, followed by an increase in HC theta power that emerges during delay 1 and persists into the presentation of the test sequence. AMY showed the most delayed response in theta power, with increases occurring during the presentation of the test sequence. The presence of theta activity in all ROIs and the difference in timing between ROIs point to theta circuitry between OFC, HC, and AMY during the encoding and maintenance of sequential information.

**Figure 2.**
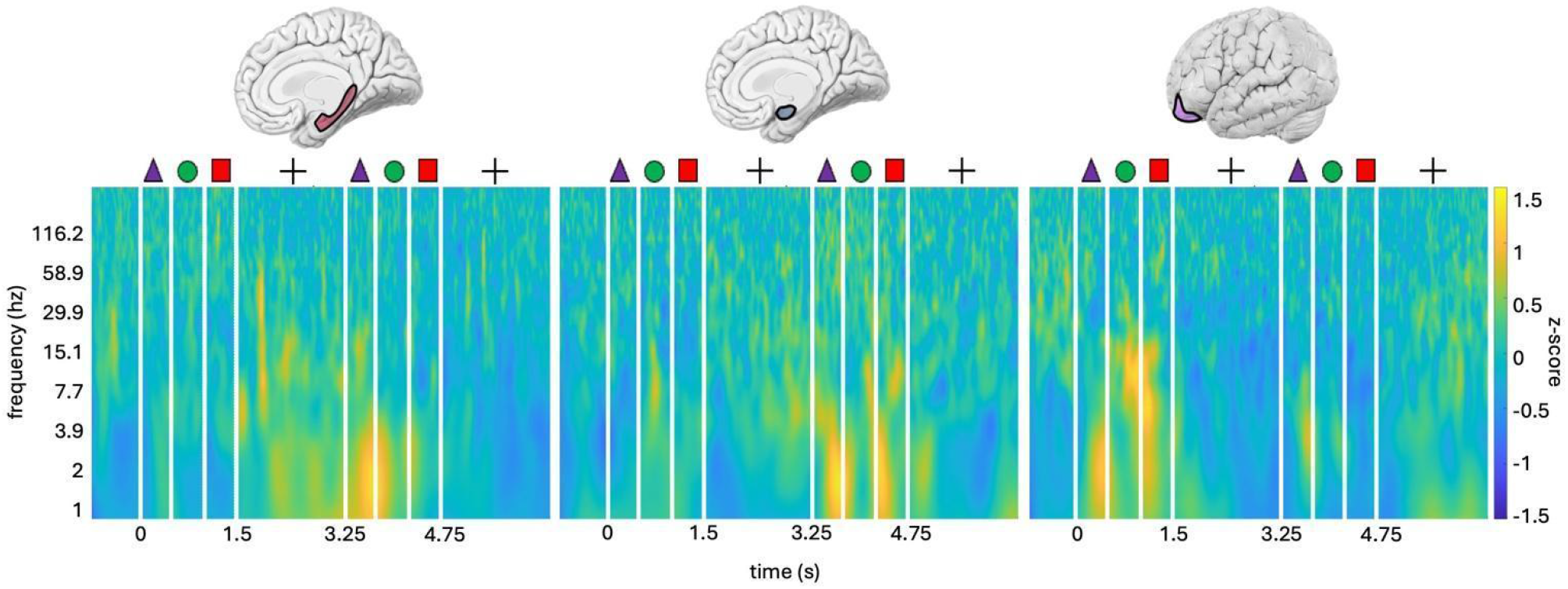
Time-frequency power during sequential WM. Theta power increases during delay 1 (1.50-3.25 s from sample onset) and the test sequence (3.25-4.75 s) in medial temporal regions (left and middle), and during the sample sequence (0-1.50 s) in OFC (right). Sample stimuli shown at the top of each figure were presented for the first 250 ms of each 500 ms stimulus presentation segment, separated by central fixation (not shown). White vertical lines represent the onset of each stimulus and delay.

### Slow theta oscillations slow down with cognitive demand in HC and OFC

To characterize slow and fast theta oscillations, we used IRASA^62^ to identify peak slow and fast theta frequencies for each channel in each ROI, using 4.5 Hz as the threshold between slow and fast theta if a single peak was identified^47^ (Figure S2). Sample peaks were computed over the presentation of the sample sequence and delay 1, and test peaks were computed over the presentation of the test sequence and delay 2 (see Figure 1A for task design). We used Wilcoxon sign rank tests to compare peak frequencies between sample and test phases.

During the test phase, participants were required to maintain both the sample sequence and the test sequence to compare the two sequences and subsequently execute a goal directed behavior. Thus, the cognitive demand during the test sequence and delay 2 was higher than during the sample and delay 1. Previous literature suggests that theta oscillations slow down with increased WM load, as a purported mechanism for more gamma cycles carrying information to fit within the theta cycle^23,24,26^. We thus operationalized the slowing of oscillations from sample to test as an indicator of functional relevance in our WM task. We hypothesized that HC slow theta would slow from sample to test, implicating HC slow theta in the maintenance and comparison of sequential information. We first tested this hypothesis in HC slow theta, and then systematically replicated analyses in AMY, OFC, and fast theta in all ROIs, to define broader theta circuitry during the maintenance and comparison of sequential information.

We reveal a slowing of slow theta oscillations from sample to test in HC (*p*=0.01, sample n=40 channels exhibiting slow theta peaks, test n=41) and OFC (*p*=0.03, sample n=21, test n=18; Figure 3, top). Slow theta frequencies did not change from sample to test in AMY (*p*=0.83, sample and test n=36). There were no significant changes in fast theta frequencies in HC (*p*=0.83 sample and test n=34), OFC (*p*=0.06, sample n=19, test n=21), or AMY (*p*=0.55, sample n=26, test n=24; Figure 3, bottom). Taken together, these results demonstrate a unique functional role of slow theta and suggest that HC and OFC interact via slow theta oscillations to maintain and process sequential information in WM.

**Figure 3.**
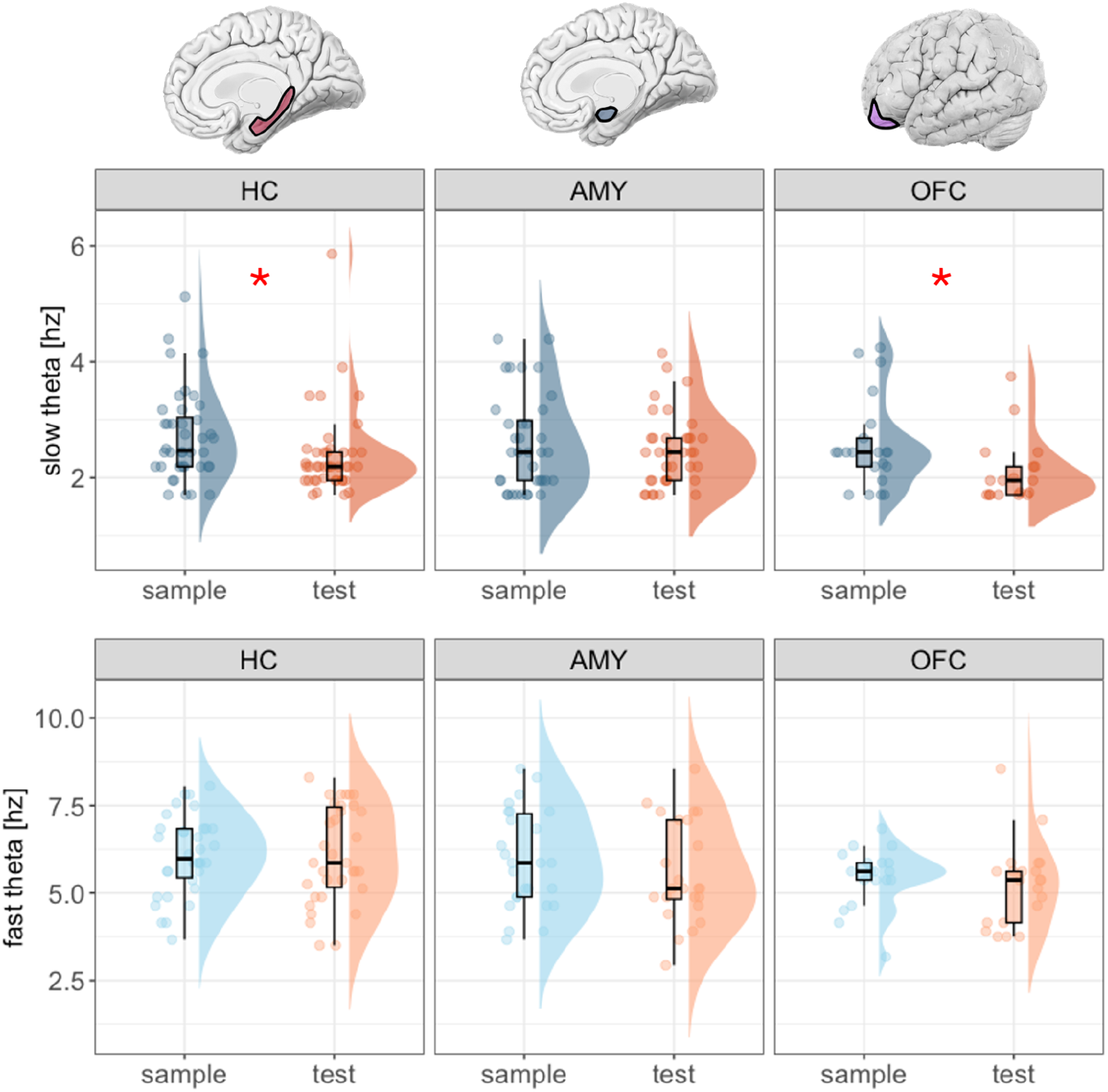
Slow and fast theta frequencies. Slow theta frequencies (top) decrease from sample to test in HC and OFC. Data are represented as individual channel datapoints, with condition probability densities and medians calculated across channels. Boxplots display medians and interquartile ranges (IQR), with whiskers extending 1.5 × IQR from the quartile. *FDR-corrected p<0.05.

### Slow and fast theta oscillations synchronize HC, AMY, and OFC

We then assessed interregional theta synchrony during sequence maintenance and comparison by computing phase locking values (PLV) during delays 1 and 2. For each interregional combination (HC-AMY, HC-OFC, AMY-OFC), PLV was computed between all within-hemisphere channel pairs at each channel’s slow theta peak, and separately at each channel’s fast theta peak^34,47^. Statistical significance was determined by comparing the observed PLV to permuted distributions generated by shuffling the phase angle time series and recomputing PLV. Based on our results demonstrating that slow theta oscillations slow from sample to test in both HC and OFC, we predicted that HC and OFC would be phase locked in slow theta during delay 1 (sample) and delay 2 (test).

Supporting our hypothesis, we observed significant slow theta PLV between HC and OFC during delay 1 (p<0.0001, n=28 channel pairs), as well as AMY-OFC (p<0.0001, n=47) and HC-AMY (p<0.0001, n=51) (Figure 4A, top). Additionally, we observed fast theta PLV between HC-OFC (p<0.0001, n=25), AMY-OFC (p<0.0001, n=37), and HC-AMY (p<0.0001, n=34) (Figure 4A, bottom). During delay 2, PLV remained significant between all ROIs at both slow (HC-OFC p<0.0001, n=24; AMY-OFC p<0.0001, n=33; HC-AMY p<0.0001, n=56) and fast (HC-OFC p<0.0001, n=23; AMY-OFC p<0.0001, n=28; HC-AMY p<0.0001, n=25) theta (Figure 4B). Our results reveal both slow and fast theta synchrony between HC, AMY, and OFC during sequence maintenance and comparison, demonstrating that the infrastructure for interregional communication is present in slow and fast theta circuits.

**Figure 4.**
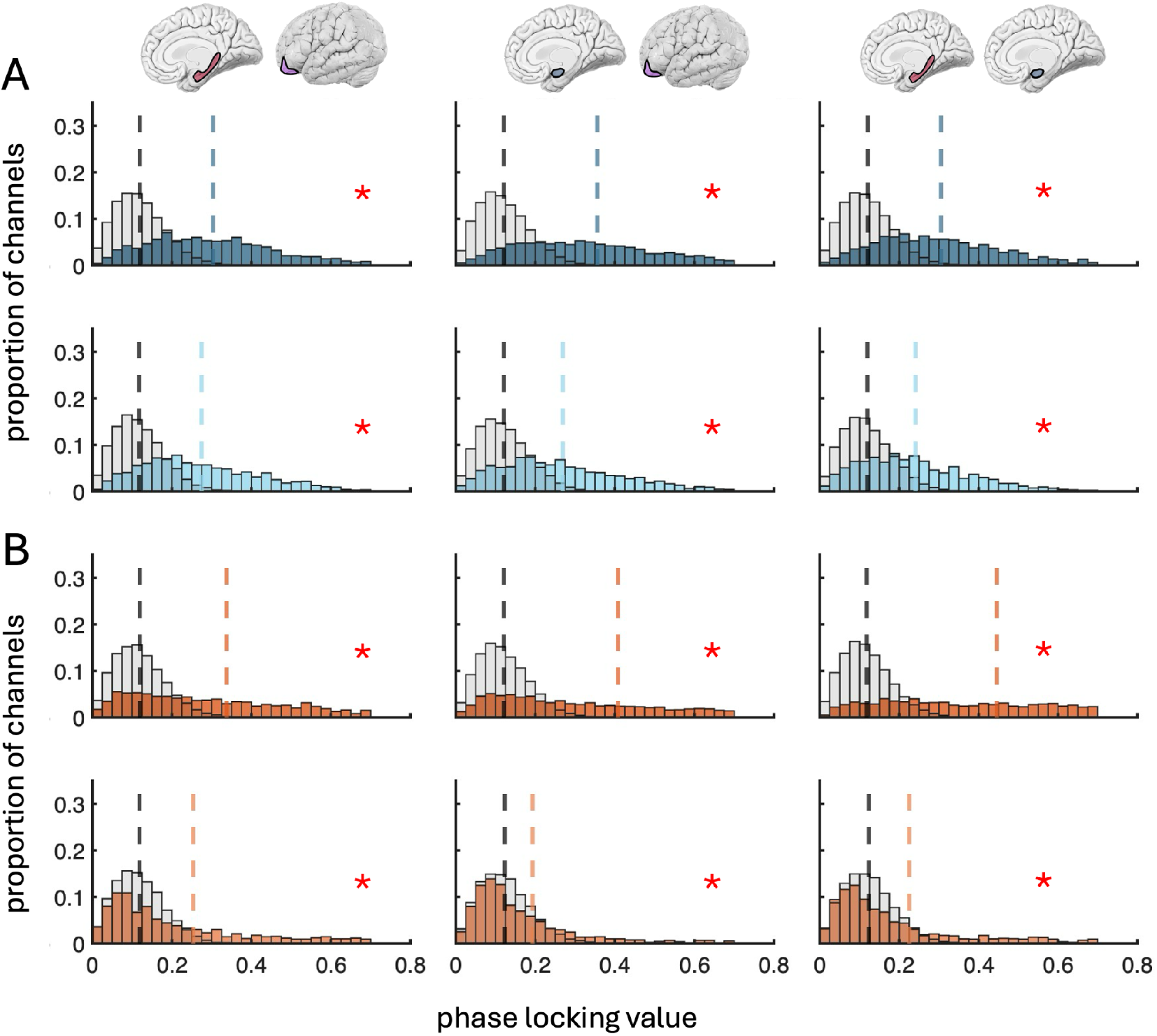
Slow and fast theta PLV during delays 1 and 2. **(A)** Significant slow (top) and fast (bottom) theta PLV in HC-OFC, AMY-OFC, and HC-AMY during delay 1. Darker colors indicate slow theta and lighter colors show fast theta for all plots. (**B**) Significant slow (top) and fast (bottom) theta PLV in HC-OFC, AMY-OFC, and HC-AMY during delay 1 and 2, same conventions as (A). Dashed vertical lines represent the distribution means. *FDR-corrected p<0.05.

### Slow and fast theta synchrony between HC and OFC exert opposing effects on behavior

Having demonstrated slow and fast theta PLV during delays 1 and 2, we then separately tested for relationships between PLV during each delay and behavioral AR and RT. One subject was excluded from the AR analyses due to an AR of 6 SDs below the mean (AR=0.73, see Table S1). Although no correlations between slow theta PLV and AR were significant (p > 0.05; Figure 5A), slow theta synchrony benefitted RT, as demonstrated by significant negative correlations with AMY-OFC PLV during delay 1 (rho= −0.38, *p*=0.02, CI [−0.63, −0.09]; Figure 5B top), and with HC-OFC PLV (rho=−0.70, *p*=0.001, CI [−0.86, −0.43]) and HC-AMY PLV during delay 2 (rho=−0.37, *p*=0.01, CI [−0.60, −0.08]; Figure 5B bottom). Fast theta synchrony between HC and OFC hindered performance, as measured by significant negative correlations with AR during delays 1 (rho=−0.52, *p*=0.046, CI [−0.76, −0.13]; Figure 5C top) and 2 (rho=−0.65, *p*=0.008, CI [−0.83, −0.36]; Figure 5C bottom), and significant positive correlations with RT during delays 1 (rho=0.64, *p*=0.002, CI [0.31, 0.83]; Figure 5D top) and 2 (rho=0.72, *p*=0.001, CI [0.48, 0.84]; Figure 5D bottom).

**Figure 5.**
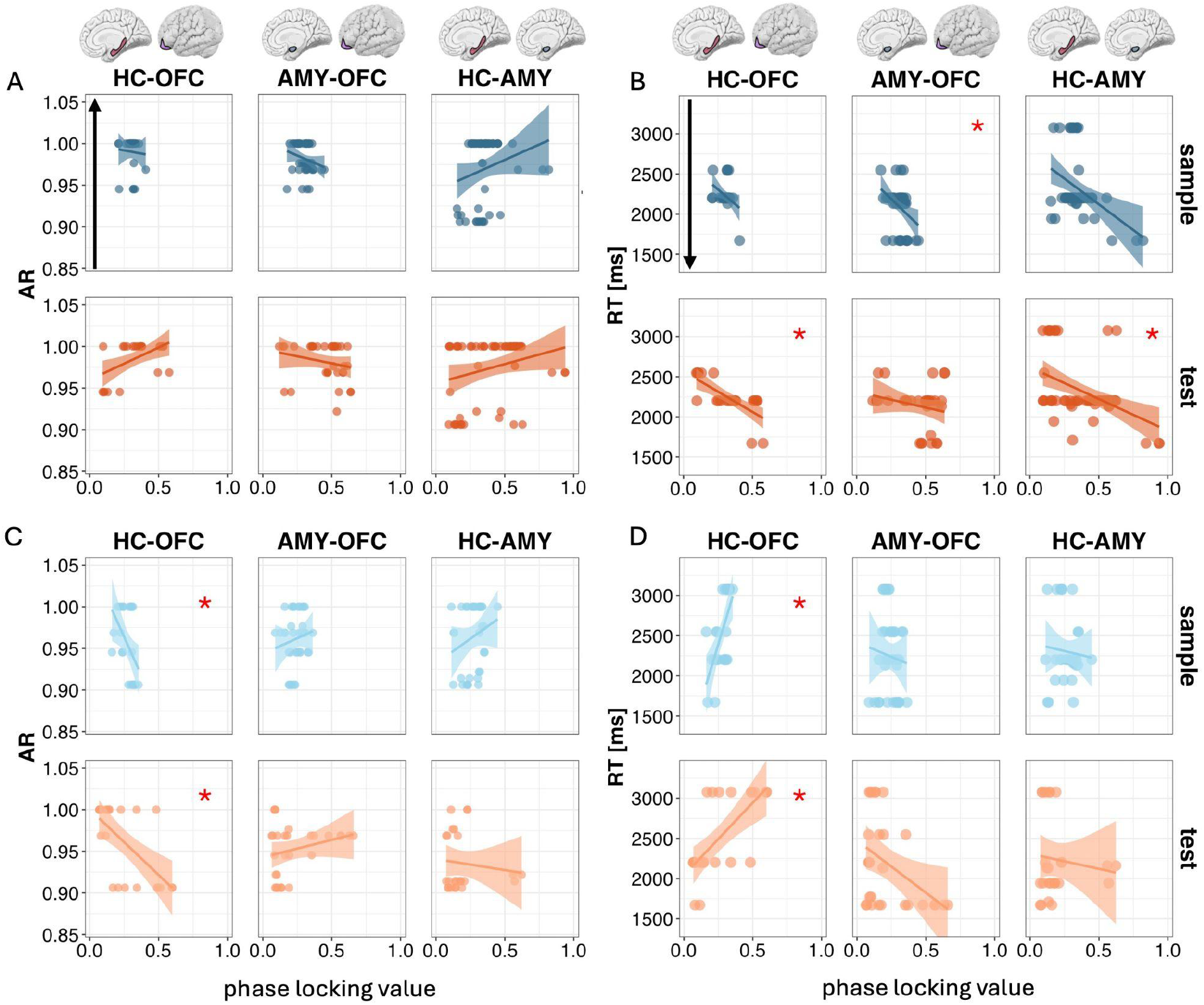
Correlations between slow and fast theta PLV during delays 1 and 2 and behavior. **(A)** Spearman’s correlation between AR slow theta PLV during delay 1 (top) and delay 2 (bottom theta PLV during delay 1. Data points represent individual channel combinations. Shading indicates the 95% confidence interval of the linear regression. Reported statistics are Spearman’s rho with bootstrapped 95% confidence intervals. The black arrow represents the direction of better performance. *FDR-corrected p<0.05. **(B)** Spearman’s correlation between RT and slow theta PLV during delay 1 (top) and delay 2 (bottom), same conventions as (A). Slow theta AMY-OFC PLV negatively correlates with RT during delay 1. Slow theta HC-OFC and HC-AMY PLV negatively correlates with RT during delay 2. **(C)** Same as (A) for fast theta. Fast theta HC-OFC PLV negatively correlates with AR during delay 1 and delay 2. **(D)** Same as (B) for fast theta. Fast theta HC-OFC PLV positively correlates with RT during delay 1 and delay 2.

Collectively, these results reveal that slow theta synchrony is associated with faster responses, with AMY-OFC effects during sequence maintenance, and HC-OFC and HC-AMY effects during sequence comparison. In contrast, fast theta HC-OFC synchrony consistently impaired both accuracy and response speed across maintenance and comparison. These results support our hypothesis that slow theta synchronization between HC and OFC supports sequential WM, perhaps due to the longer temporal window within a cycle for single item representations^23^, while fast theta impaired performance due to its shorter cycle length.

### High-frequency broadband activity is coupled to slow theta oscillations in HC and OFC during sequence maintenance and comparison

We then assessed slow and fast theta-HFB PAC within our three ROIs during sequence maintenance by computing the mean vector length (MVL)^35^ in delays 1 and 2, using the same individual peak theta frequencies as in PLV analyses. Statistical significance was assessed by comparing the observed MVL to permuted distributions generated by shuffling the phase angle time series and recomputing MVL. Based on literature positing slow theta as the optimal frequency for organizing multiple item representations in WM^23,24^, we predicted that HFB would be coupled to slow theta oscillations in both HC and OFC, which would reinforce the proposal that slow theta provides optimal temporal windows for organizing sequential information within these regions.

Supporting our hypothesis, we observed significant slow theta-HFB MVL in HC (*p*=0.0008), OFC (*p*=0.0007), and AMY (*p*=0.0007) during delay 1 (Figure 6A). We also observed significant fast theta-HFB PAC in HC (*p*=0.039; Figure 6B), demonstrating both slow and fast theta PAC in HC during sequence maintenance. By contrast, fast theta-HFB PAC was not significant in AMY (*p*=0.056) or OFC (*p*=0.57). Results reveal that slow theta oscillations modulate HFB activity in all three regions during the maintenance of sequential information.

**Figure 6.**
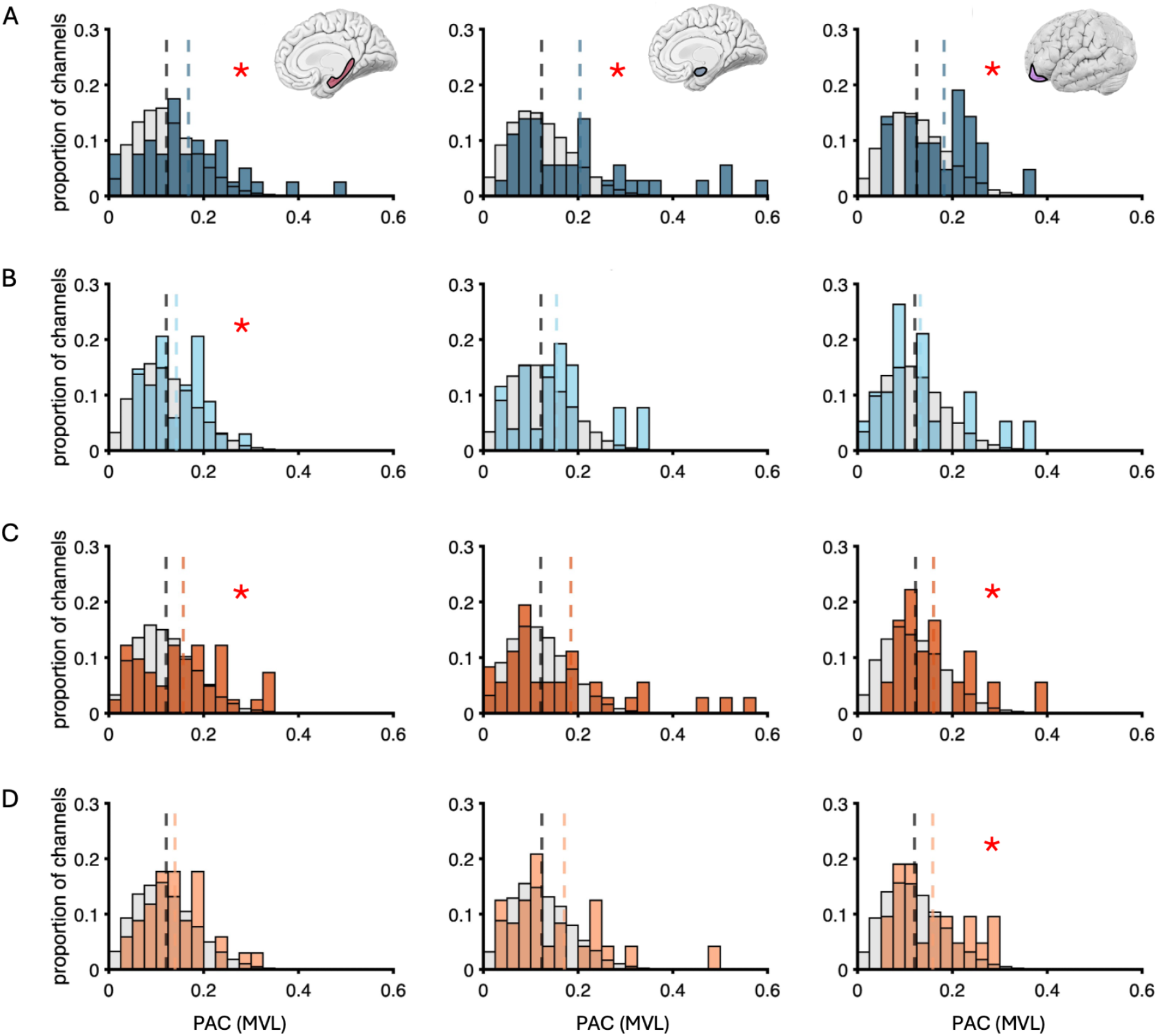
Slow and fast theta PAC MVL during delays 1 and 2. (**A**) Significant slow theta MVL (top) in HC, AMY, and OFC during delay 1. *FDR-corrected p<0.05. (**B**) Significant fast theta MVL in HC during delay 1, same conventions as (A). (**C**) Same as (A) for delay 2. Significant slow theta MVL in HC and OFC. (**D**) Same as (B) for delay 2. Significant fast theta MVL in OFC. Dashed vertical lines represent the means of the distributions.

During delay 2, slow theta MVL remained significant in HC (p<0.007) and OFC (p<0.04), but was not significant in AMY (*p*=0.16; Figure 6C). We also observed significant fast theta-HFB PAC in OFC (*p*=0.03) (Figure 6D), demonstrating both slow and fast theta PAC in OFC during sequence comparison. Fast theta-HFB PAC was not significant in HC (*p*=0.08) or AMY (*p*=0.16). No correlations between PAC during delay 1 or 2 and AR or RT were significant (*p*≥0.05).

### Initial evidence that single neurons encode sequential order through slow theta phase coding across the HC-OFC circuit

Finally, we analysed slow and fast theta phase coding in the subset of 10 patients with microwire implants. Rayleigh’s test revealed significant slow theta phase coding for HC neurons preferring stimulus 2 and OFC neurons preferring stimulus 3, with no significant fast theta phase coding in any region. Given sample size limitations, these findings are preliminary and presented in full in Supplemental Text S3 and Figure S3.

## Discussion

Our findings establish that HC and OFC form a slow theta circuit for sequential WM, with dissociable roles of slow and fast theta synchrony within the same network. When synchronized across HC and OFC, slow theta facilitated response speed, whereas fast theta synchrony between HC and OFC consistently impaired response accuracy and speed (Figure 7). This frequency dissociation was present at the local level, with persistent slow theta-HFB PAC in HC and OFC through sequence maintenance and comparison, corroborating a complementary intraregional mechanism for organizing information in WM that has been well-documented across tasks and species^25,35,42–44^. By contrast, fast theta-HFB PAC was limited to HC during sequence maintenance and OFC during sequence comparison. Together, these findings reveal a frequency-opponent mechanism in which the same circuit can facilitate or disrupt sequential WM depending on frequency, with fast theta synchrony being detrimental to subsequent behavior.

**Figure 7.**
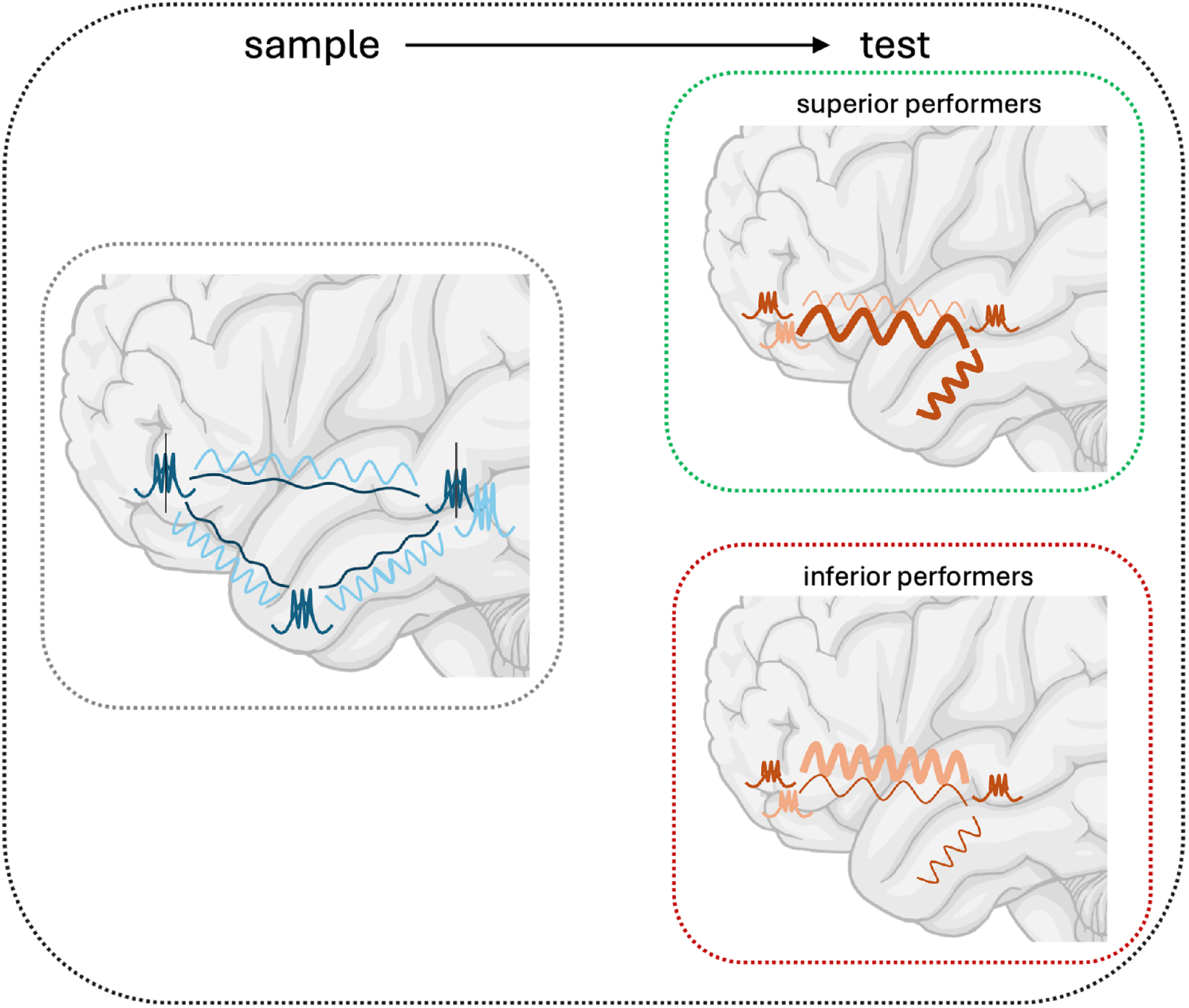
Schematic of main findings. Group level analyses revealed significant slow and fast theta PLV between all regions across both sample (left) and test (right, not shown) phases of the task. Slow theta-HFB PAC was present in all three regions during sample, with sustained PAC in HC and OFC during test. Slow theta phase coding of sequence neurons was observed in HC and OFC (see S3). Fast theta-HFB PAC was present in HC during sample and shifted to OFC during test. Slow and fast theta PLV between HC and OFC differentially explained behavioral performance during test, with slow theta linked to faster RTs and fast theta hindering accuracy and RTs. Blue represents sample slow (dark blue) and fast (light blue) theta. Orange represents test slow (dark orange) and fast (light orange) theta. Thicker lines signify stronger synchrony in superior (top) or inferior (bottom) performers.

Prior to testing hypotheses, we removed pathological channels and trials to ensure generalizability of findings to healthy populations. Behavioral performance in our neurosurgical patients was comparable to that of healthy controls^46,47^, validating our findings as representative of typical WM function. We then established theta activity across all three regions, each with a qualitatively distinct temporal profile: OFC theta power peaked during sequence encoding, HC theta emerged during maintenance and persisted into the test sequence, and AMY theta peaked during the test sequence. We then extended previous reports of slow and fast theta sub-bands in HC^48–51^ and cortex^47^ by demonstrating that both theta oscillations are present across all three regions in our network. We then revealed a functional role of theta oscillations during the WM-DMS task, revealing a slowing of slow theta from sample to test in HC and OFC and further suggesting that these two regions interact during WM of sequential information. These initial tests demonstrated the functional role of slow theta, the oscillation we hypothesized would support sequential WM, and further suggested similar neural mechanisms of sequential WM in HC and OFC. These findings are consistent with literature suggesting that increased cycle length of theta oscillations, or theta slowing, provides the architecture for single items to be represented in WM^21–24^. The slowing of slow theta and lack thereof in fast theta from sample to test also extends previous findings of functionally distinct slow and fast theta oscillations in long term memory processes to WM processes^47,49,52^.

Our PLV results demonstrate sustained slow and fast theta synchrony spanning the HC-OFC circuit throughout the WM task, establishing the infrastructure for multiplexed interregional communication^42,43,63,64^. The frequency of this synchrony impacted WM performance, with slow theta synchrony predicting faster RTs across multiple circuit connections and HC (HC-OFC and HC-AMY) effects emerging during sequence comparison. By contrast, fast theta HC-OFC synchrony consistently increased RT and decreased accuracy across maintenance and comparison. This aligns with previous theoretical and empirical studies suggesting that slow theta cycles provide optimal temporal windows for organizing and transferring sequential information, while fast theta cycles may disrupt these processes^23–26,45^. We speculate that HC-OFC slow theta synchrony is most critical for processing efficiency during high cognitive demand. Additionally, AMY showed demand-dependent slow theta behavioral relationships, with AMY-OFC synchrony explaining faster RTs during maintenance and HC-AMY synchrony explaining faster RTs during comparison. This pattern is consistent with AMY’s delayed theta onset and maintenance-specific PAC, suggesting that AMY dynamically modulates response speed through different anatomical pathways depending on cognitive demand. These results corroborate fMRI findings demonstrating increased delay-period HC-frontal connectivity with increased visual WM load^65^ and AMY modulation of response speed^14^, providing mechanistic insight into how these regions coordinate to support sequential WM.

Our MVL results demonstrate slow theta PAC in HC, AMY, and OFC during the maintenance period of the task, revealing the infrastructure for slow theta oscillations to modulate HFB activity and maintain sequential information, and extending theoretical models of theta-gamma neural coding in WM^23,27^ to theta-HFB PAC in the human brain. Slow theta-HFB PAC was sustained during comparison within HC and OFC, aligning with our PLV findings and supporting the hypothesis that these regions form a coordinated slow theta circuit. These results align with studies demonstrating theta-HFB PAC in HC and AMY during WM maintenance^25^, and extend these findings to OFC. In contrast, fast theta PAC was only present in HC during maintenance and OFC during comparison, suggesting that fast theta PAC may serve region-specific roles that, when synchronized across the circuit, interfere with the more optimal slow theta organization.

These findings demonstrate distinct roles of slow and fast theta synchrony across frontotemporal circuits in sequential WM. We overcame the limitations of spatial sampling common in invasive recordings^66^ by densely sampling ROIs with sEEG macroelectrodes. All subjects had epilepsy, which can limit generalizability despite strict electrode selection excluding pathological tissue^47,66^. Critically, however, patients in our sample demonstrated intact behavioral performance comparable to healthy controls, suggesting that the mechanisms we identified reflect typical WM function^46^. Our synchrony-behavior relationships are correlational and cannot establish causality, but are strongly supported by previous studies demonstrating that slow theta tACS benefits temporal memory while fast theta tACS does not^53–55^. To further investigate the spatial specificity of such results, future studies could apply invasive direct electrical stimulation to OFC, HC, and AMY to perturb slow and fast theta oscillations and observe the behavioral effects^67^. To further investigate the therapeutic efficacy of such interventions, future clinical trials could be targeted to patients with memory deficits.

In conclusion, our findings reveal that within the same HC-OFC circuit, slow theta oscillations facilitate sequential WM through complementary slowing, interregional synchrony, and sustained PAC, while fast theta oscillations disrupt performance, potentially through competing dynamics that interfere with sequential information transfer. This dual frequency-opponent mechanism establishes a framework where the same circuit architecture can produce opposing cognitive outcomes depending on the theta frequency at which the circuit is synchronized. These findings advance our understanding of theta oscillations in human cognition and identify potential frequency- and anatomically-selective therapeutic targets for WM deficits.

## Methods

### Human subjects

Subjects were 21 neurosurgical patients (7 females; 13-54 years of age; M ± SD, 30 ± 11.2 years) undergoing stereotactic EEG (sEEG) monitoring as part of clinical management of seizures. Microwire recordings were obtained from 10 of these subjects (2 females; 21-54 years of age; M ± SD, 32 ± 9.5 years). Demographic details are provided in Table S1. Subjects were selected from a larger pool based on non-pathological coverage of hippocampus (HC), amygdala (AMY), and/or orbitofrontal cortex (OFC; i.e., electrodes localized to HC, AMY, and/or OFC and outsize seizure onset zones). Patients were recruited from the University of California, Irvine, University of California, Davis, Icahn School of Medicine at Mount Sinai, University of California, San Francisco, UCSF Benioff Children’s Hospital, and Nationwide Children’s Hospital. Written informed consent was obtained from subjects aged 18 years and older and from the guardians of subjects younger than 18 years, and written assent was obtained from subjects aged 13-17 years. The Institutional Review Board of the institution at which the patient was enrolled approved all procedures in accordance with the Declaration of Helsinki.

### Task design

Subjects completed a working memory delayed match-to-sample (WM-DMS) task where each trial presented a sequence of three shape stimuli (sample sequence; Figure 1A)^46,57,58^. The stimuli varied in identity (i.e., shape and color), and each was presented in one of four spatial locations on the screen (top, bottom, left, or right) for 250ms, followed by a 250ms interstimulus interval. After the last shape, there was a 2-second delay (delay 1) where subjects were to maintain the sample sequence. Proceeding delay 1, a second sequence of three shapes was presented (test sequence). The test sequence was either an exact match to the sample sequence, or a mismatch in one of three ways: temporal order, spatial location, or identity. After the last shape, there was a second 2-second delay (delay 2), after which subjects indicated whether the test sequence was a match or a mismatch to the sample sequence. The response was self-paced. The central fixation crosshair remained on screen for the duration of the task. Following eight practice trials, subjects completed 128 trials with a break every 16 trials. The four test conditions were evenly counterbalanced and presented in pseudo-randomized order with four trials per condition every 16 trials. For half of all trials, a star flashed for 100ms midway through both delays at randomly jittered times ranging from 1000-1150ms from the onset of the delay. The star trials were evenly counterbalanced across conditions and appeared on eight of every 16 trials. The other half of trials were considered control trials and analyzed in this study. Behavioral data were collected using custom-built MATLAB (MathWorks Inc., Natick, MA) scripts with the Psychtoolbox-3 software extension.

Behavioral accuracy rate (AR) was calculated per subject as the hit rate (i.e., proportion of match sequences that were correctly identified as match) minus false alarm rate (proportion of mismatched sequences that were incorrectly identified as match^68^. This procedure defines chance accuracy as zero and corrects for differences in trial counts between conditions as well as an individual’s tendency to respond match/mismatch. Response time (RT) was calculated per subject across all correct, no-star trials.

### Electrode placement and localization

Platinum macroelectrodes (5-10-mm intercontact distance, 4-mm diameter) were surgically implanted for extra-operative sEEG recording based solely on the clinical needs of each patient. Three-dimensional electrode reconstructions were created by co-registering post-implantation planar x-ray images of the cortical surface with preoperative T1-weighted spoiled gradient-echo magnetic resonance (MR) images, as implemented in FieldTrip^69^. Electrodes were localized according to individual anatomy and transformed into standard MNI space for visual representation across subjects (Figure 1B). Benke-Fried microelectrodes were included in the clinical implant for 10 of the subjects in this study. Each Benke-Fried electrode contained eight micro-wires attached to the deepest contact on the sEEG macroelectrode. Micro-wire electrodes were localized to the region of the sEEG contact to which they were attached.

### Data acquisition and preprocessing

sEEG data were acquired using Nihon Kohden and Natus recording systems at a minimum sampling rate of 1kHz, and data sampled >1kHz were resampled to 1kHz offline. Raw electrophysiology data were bandpass filtered from 0.1-300 Hz using finite impulse response filters and 60-Hz line noise harmonics were removed using discrete Fourier Transform. Continuous study data were demeaned, epoched into 8.5-s trials (−1 s from the onset of the sample sequence to +1 s from the offset of delay 2), and manually inspected blind to electrode locations and experimental task parameters. Electrodes overlying seizure onset zones^70^ and electrodes and epochs displaying epileptiform activity or artifactual signal (from poor contact, machine noise, etc.) were excluded. The remaining electrodes were re-referenced to neighboring electrodes within the same anatomical structure using consistent conventions (i.e., deep – surface)^34,59^. A channel was discarded if it did not have an adjacent neighbor, its neighbor was in a different anatomical structure, or both it and its neighbor were in white matter. Bipolar montages were used to minimize contamination from volume conduction prior to analysis of functional connectivity^71,72^. Data were manually re-inspected to reject any trials with residual noise, and error trials were excluded^34,59^.

Microwire data were acquired simultaneously with the sEEG data using Nihon Kohden recording systems at a sampling rate of 32kHz. If the macroelectrode that the microwires were attached to had been removed in the sEEG preprocessing pipeline, all eight microwire channels attached to the tip of that probe were excluded. The same trials that were removed in the sEEG data were also removed in the microwire data. The microwire data were separately bandpass filtered from 0.1-300 Hz for analysis of local field potentials (LFPs) and 300-3kHz for analysis of neuronal spikes, both described below. LFP data were re-referenced using a leave-one-out common average scheme. For each of eight target channels per probe, signals from the other seven channels were averaged and subtracted from the target channel.

All results are based on analysis of non-pathologic, artifact-free electrodes in regions of interest (ROIs), ensuring that data represent healthy tissue^73^. On average, 53 (SD: 6) seizure- and artifact-free, correct trials were analyzed for each subject, with an average of 53 (SD: 5) trials analyzed in subjects with both sEEG and microwire data. Preprocessing routines utilized functions from the FieldTrip toolbox for MATLAB^74^.

### sEEG signal processing

#### Power

We first subtracted the event-related potential (ERP) from each trial. We then zero-padded the data segments to 15 s and calculated the time-frequency spectrum. We extracted narrowband time series from 40 logarithmically spaced frequencies between 1 and 200 Hz using convolution of broadband data with complex Morlet wavelets^34,59,75^, with standard deviations of 40 logarithmically spaced values from 2 to 24 Hz. This effectively created bandpass filters with pass bands ranging from 2 to 24 Hz around the center frequencies. The resulting complex analytic signal was converted to power by taking the absolute value of the squared complex values. These power outputs were epoched −0.75 to 6.5 s from sample onset (with the offset of delay 2 at 6.5 s) and z-scored on null distributions using statistical bootstrapping. For each electrode and frequency, N random samples (N = number of trials in that subject’s dataset) were selected from the vectorized time series data across all trials and averaged. This process was repeated 1000 times to create normal distributions. Raw power data were then z-scored on the bootstrapped distributions^34,44,59–61^.

#### Oscillatory peak detection

We used irregular-resampling auto-spectral analysis (IRASA) to separate true oscillatory components from the aperiodic 1/*f* slope, as implemented in FieldTrip^62,69^. Study data segments were epoched 0 to 3.25 s (sample sequence and delay 1) and 3.25 to 6 s (test sequence and delay 2), zero-padded to the next power of 2, and analyzed from 1 to 50 Hz. The IRASA method compresses and expands the epoched data with non-integer resampling factors to redistribute oscillatory components while leaving the 1/*f* distribution intact. For each original and resampled data trace, the auto-spectrum was calculated using the fast Fourier transform^76^ after applying a Hanning window. The median was taken from the resampled auto-spectra to obtain the 1/*f* component for each electrode and subtracted from the original power spectrum to isolate oscillatory residuals^62,69^. Peak detection was performed on the oscillatory residuals within the theta range (1.5–9 Hz) using a minimum prominence threshold of 0.5, and a threshold of 4.5 Hz was used to distinguish slow from fast theta. When a single peak was detected, it was assigned to the appropriate sub-band based on its frequency. When two peaks were detected, the lower was assigned to slow theta and the higher to fast theta. When more than two peaks were detected within a sub-band, the peak with the greatest prominence was retained^47^. See Figure S2 for two examples of detected slow and fast theta peaks.

#### Phase-amplitude coupling

Phase-amplitude coupling (PAC) was measured using the mean vector length (MVL) between slow and fast theta phase, respectively, and high-frequency broadband (HFB) amplitude following removal of the ERP^47,61^. We used slow and fast theta peak frequencies for each channel, as determined by oscillatory peak detection. Narrowband theta time series were extracted using Morlet wavelets as described above for time-frequency power, with a standard deviation of 3 Hz. Phase angles were converted using MATLAB’s built-in angle function. To construct HFB timeseries, we used Morlet wavelet convolution at 40 linearly spaced frequencies between 75 and 145 Hz, with standard deviations increasing linearly from 10 to 20 Hz to create pass bands ranging from 10 to 20 Hz around the center frequency. The resulting narrowband time series were first converted to power and z-score normalized within each frequency^34^. The mean z-scored power was then taken across frequencies at each time point. We separately measured the phase at which peak HFB power occurred during delays 1 and 2 for both slow and fast theta phases on a per trial basis. This procedure yielded one slow and one fast theta phase angle per delay per trial. These angles were then used to compute the MVL across trials per delay.

Null distributions of PAC were simulated using a permutation technique where we vectorized the phase angle data from all trials within each channel and subsequently circularly rotated the data around a randomly chosen time point from within the middle 80% of the timeseries. We recomputed the MVL on the circularly rotated data. We repeated this procedure 1000 times to generate normal distributions. This procedure shuffles the timing of the amplitude envelope relative to the phase without altering the autocorrelation structure in either time series, thus eliminating differences in input data that could confound PAC^77^.

#### Phase-locking value

Theta phase synchrony between ROIs was measured using the phase-locking value (PLV)^64^ following removal of the ERP^47,59,61^. Slow and fast theta PLV were computed separately, using the same peak frequencies and phase angle extraction methods described above for PAC^34,47,78^. Instantaneous slow and fast phase angles were separately extracted from delays 1 and 2 on a per trial basis, and PLV was calculated for each within-hemisphere interregional electrode pair. This method calculates the consistency in electrode-pair phase differences across timepoints per trial, yielding one slow and one fast theta PLV per delay per trial. Null distributions of PLV were simulated using the permutation technique described above for PAC^61^. This procedure shuffles the timing of the phase of one electrode without altering the original data and thus eliminates differences in input data that could confound PLV.

### Microwire signal processing

#### Spike detection

Neuronal spikes were detected as threshold crossings^79^ five standard deviations above or below the mean of the entire time series. Spikes that occurred within 16 time points of each other were not included, eliminating spikes that occur within 0.5 ms of each other due to limitations of effectively clustering spikes that close in time^80^. Spikes were removed if they occurred within 32 timepoints (1 ms) of the beginning or end of the task recording due to the inability to capture the full waveform. For each waveform peak detected, 32 timepoints (1 ms) on either side of the detected peak were extracted as the spike waveform, from which waveform maxima were calculated. If a waveform had more than one threshold crossing, it was aligned to the first peak detected. We further excluded waveforms if the sign of the first derivative did not change from the first peak to the second peak because this would indicate noise in the extracted waveform.

#### Spike sorting

Microwire channels with fewer than 400 detected waveforms across the entire duration of the task were excluded from further analyses due to the inability to generate template waveforms using clustering^81^. For channels with more than 50K waveforms, 50K were selected at random for runtime optimization and were later used to generate template waveforms. All remaining waveforms were used for spike sorting (i.e., 400-50K total waveforms per channel). Waveforms with a peak above zero were classified as up going waveforms and waveforms with a peak below zero were classified as down going. A HAAR covariance matrix was created using four HAAR coefficients^82,83^. Up and down going waveforms were either clustered into noise or “real clusters” in HAAR space^82,83^ using DBScan, a density-based clustering algorithm with a dynamic epsilon selection, as implemented in MATLAB^81,84^. Clusters were then z-scored across time. To further remove outlying waveforms in each cluster that deviated from the rest of the waveforms on a cluster, we removed any waveform that deviated by a z-score of ±3 at any point in time and sorted it as noise. This procedure accounts for any misclustered waveforms. Mean waveforms were calculated by taking the average within each remaining real cluster and used for template matching.

For template matching, each waveform detected on the channel was assigned to a new cluster by computing the Euclidean distance between each waveform on the channel and each template waveform. Waveforms were assigned to the template with the closest Euclidean distance. All waveforms assigned to the same template were then considered a cluster. We used the difference between the distance to the closest template and the distance to the second closest template as a measure of discriminability for each waveform^79^. Waveforms that were well sorted into a cluster would be very close to one template and far from even the second closest template, yielding high discriminability. We then found the inflection point of the density distribution of all waveforms for that cluster as a function of discriminability, and sorted all waveforms that had a discriminability index before the inflection point as noise. If there was more than 1 inflection point, the ratio of noise to real waveforms as determined by the DBScan clustering algorithm was assumed, and that same proportion of waveforms with the lowest discriminability index was sorted into noise. A final trim was done on each real cluster by excluding the most deviant 1% of waveforms in each cluster. Clusters with fewer than 100 waveforms after the final trimming were considered to have a firing rate too low for further analysis and excluded. All remaining clusters were then manually inspected to reject any with residual noise^79^. Single units were determined by the neuronal clusters output.

#### Neuronal stimulus preference of sample sequence during delay 1

For each unit, the average firing rate during the presentation of each stimulus and ISI (500ms total) was calculated across all trials. Stimulus preference was determined per unit as the stimulus with the maximal firing rate^37^. Units with no firing during stimulus presentation or equal firing for two or more stimuli were excluded from further analyses because they did not exhibit neuronal stimulus preference. Units included in further analyses were required to have a firing rate above an average of 0.5 Hz per trial, and a minimum of one spike during stimulus presentation and one spike during delay 1.

#### Phase coding by neuronal stimulus preference

Neuronal phase coding was measured by assessing the consistency in phase across spike times for each unit exhibiting neuronal stimulus preference. Equivalent to the methods stated above for PAC and PLV, we used slow and fast theta peak frequencies for each microwire channel, as determined by oscillatory peak detection. Using the unit spike times detected for each neuron, we assessed the instantaneous slow theta phase at all unit spike times per trial during delay 1. We then grouped neurons by the stimulus preference determined by unit activity during the presentation of the sample sequence, and pooled all phases at spike times across neurons within each category for statistical assessment of phase uniformity^40,85^. P-values were FDR corrected for the 3 stimulus preference categories tested for a given condition in MATLAB using the mafdr function. In the case where there were only two stimulus preferences present in the neurons, p-values were FDR corrected for two categories.

### Statistical analyses

#### Neurophysiology

Changes in slow and fast theta frequencies from sample to test were assessed using paired Wilcoxon signed rank tests, as implemented in *R*^*86*^. MVL was assessed using unpaired Mann-Whitney U rank sum tests to compare the observed raw MVL values to the permuted (null) distributions separately for delay 1 (sample) and 2 (test). The same tests applied to MVL were applied to PLV. PLV data were averaged over trials to obtain one datapoint per channel, matching the format of MVL. Changes in MVL from sample to test were assessed using paired Wilcoxon signed rank tests and changes in the theta phase at peak HFB were assessed using the circular equivalent^85^. All *p*-values were FDR-corrected across the three ROIs, slow and fast theta frequencies, and sample and test conditions, correcting for a total of 12 comparisons.

#### Brain-behavior relationships

Relationships between MVL and AR were assessed using Spearman’s correlation, with AR as a repeated measure across channels, as implemented in *R*^86,87^. Correlations were computed separately for each region, frequency band, and condition. Relationships between MVL and RT were also assessed using Spearman’s correlation separately for each region, frequency band, and condition. RTs were averaged over trials to obtain one RT for each subject, matching the format of AR, and subsequently used as a repeated measure across channels. The same tests applied to MVL were applied to PLV, with PLV data averaged over trials and computed separately for each frequency band, condition, and region. All correlations were computed at the channel pair level, with behavioral measures treated as repeated across channel pairs within each subject. 95% confidence intervals for Spearman’s correlation were estimated using percentile bootstrapping with 1000 resampling iterations, as implemented R. All *p*-values were FDR-corrected for 12 comparisons, corresponding to two frequency bands (slow and fast theta), two conditions (sample and test), and three regions (HC, AMY, and OFC or HC-AMY, HC-OFC, and AMY-OFC).

#### Phase coding

Spikes from neurons were grouped by stimulus preference, as described above, and statistically tested for clustering by theta phase using Rayleigh’s test of uniformity^85^. P-values were FDR-corrected across the three stimulus preferences. We then tested for phase clustering differences between the three groups of neurons exhibiting stimulus preference using the circular Kruskal-Wallace test^85^.

## Funding

NIH BRAIN Initiative R00NS115918, NINDS R01NS021135

## Acknowledgements

This research was supported in part through the computational resources and staff contributions provided for the Quest high performance computing facility at Northwestern University which is jointly supported by the Office of the Provost, the Office for Research, and Northwestern University Information Technology.

## Notes

### Competing Interest Statement

The authors have declared no competing interest.

## References

1. Dunning, D. L., Westgate, B. & Adlam, A.-L. R. A meta-analysis of working memory impairments in survivors of moderate-to-severe traumatic brain injury. Neuropsychology 30, 811–819 (2016).

2. Zhou, J., Li, J., Zhao, Q., Ou, P. & Zhao, W. Working memory deficits in children with schizophrenia and its mechanism, susceptibility genes, and improvement: A literature review. Front. Psychiatry 13, (2022).

3. Ortega, R. et al. Neurocognitive mechanisms underlying working memory encoding and retrieval in Attention-Deficit/Hyperactivity Disorder. Sci. Rep. 10, (2020).

4. McDowell, S., Whyte, J. & D’Esposito, M. Working memory impairments in traumatic brain injury: evidence from a dual-task paradigm. Neuropsychologia 35, 1341–1353 (1997).

5. Forbes, N. F., Carrick, L. A., McIntosh, A. M. & Lawrie, S. M. Working memory in schizophrenia: a meta-analysis. Psychol. Med. 39, 889–905 (2009).

6. Adam, K. C. S. et al. Beyond Routine Maintenance: Current Trends in Working Memory Research. J. Cogn. Neurosci. 37, 1035–1052 (2025).

7. Sternberg, S. Memory-scanning: mental processes revealed by reaction-time experiments. Am. Sci. 57, 421–457 (1969).

8. Alenazi, M. F. et al. Spatial Binding Impairments in Visual Working Memory following Temporal Lobectomy. eneuro 9, ENEURO.0278-21.2022 (2022).

9. Delogu, F., Nijboer, T. C. W. & Postma, A. Binding “When” and “Where” Impairs Temporal, but not Spatial Recall in Auditory and Visual Working Memory. Front. Psychol. 3, (2012).

10. Allen, R. J., Baddeley, A. D. & Hitch, G. J. Is the binding of visual features in working memory resource-demanding? J. Exp. Psychol. Gen. 135, 298–313 (2006).

11. Fuster, J. M. & Alexander, G. E. Neuron Activity Related to Short-Term Memory. Science 173, 652–654 (1971).

12. Goldman-Rakic, P. S. Cellular basis of working memory. Neuron 14, 477–485 (1995).

13. Jeneson, A. & Squire, L. R. Working memory, long-term memory, and medial temporal lobe function. Learn. Mem. 19, 15–25 (2012).

14. Schaefer, A. et al. Individual Differences in Amygdala Activity Predict Response Speed during Working Memory. J. Neurosci. 26, 10120–10128 (2006).

15. Zheng, J. et al. Amygdala-hippocampal dynamics during salient information processing. Nat. Commun. 8, 14413 (2017).

16. Wang, J. & Barbas, H. Specificity of Primate Amygdalar Pathways to Hippocampus. J. Neurosci. 38, 10019–10041 (2018).

17. Johnson, E. L. et al. Orbitofrontal cortex governs working memory for temporal order. Curr. Biol. 32, R410–R411 (2022).

18. Thiebaut De Schotten, M., Dell’Acqua, F., Valabregue, R. & Catani, M. Monkey to human comparative anatomy of the frontal lobe association tracts. Cortex 48, 82–96 (2012).

19. Dejerine, J. Anatomie Des Centres Nerveux: Méthodes Générales d’étude. vol. 1 (1895).

20. Tesche, C. D. & Karhu, J. Theta oscillations index human hippocampal activation during a working memory task. Proc. Natl. Acad. Sci. 97, 919–924 (2000).

21. Buzsáki, G. Theta Oscillations in the Hippocampus. Neuron 33, 325–340 (2002).

22. Jensen, O. & Tesche, C. D. Frontal theta activity in humans increases with memory load in a working memory task. Eur. J. Neurosci. 15, 1395–1399 (2002).

23. Lisman, J. E. & Jensen, O. The Theta-Gamma Neural Code. Neuron 77, 1002–1016 (2013).

24. Lisman, J. E. & Idiart, M. A. P. Storage of 7 ± 2 Short-Term Memories in Oscillatory Subcycles. Science 267, 1512–1515 (1995).

25. Daume, J. et al. Control of working memory by phase–amplitude coupling of human hippocampal neurons. Nature 629, 393–401 (2024).

26. Axmacher, N. et al. Cross-frequency coupling supports multi-item working memory in the human hippocampus. Proc. Natl. Acad. Sci. 107, 3228–3233 (2010).

27. Canolty, R. T. & Knight, R. T. The functional role of cross-frequency coupling. Trends Cogn. Sci. 14, 506–515 (2010).

28. Ray, S., Crone, N. E., Niebur, E., Franaszczuk, P. J. & Hsiao, S. S. Neural Correlates of High-Gamma Oscillations (60–200 Hz) in Macaque Local Field Potentials and Their Potential Implications in Electrocorticography. J. Neurosci. 28, 11526–11536 (2008).

29. Watson, B. O., Ding, M. & Buzsáki, G. Temporal coupling of field potentials and action potentials in the neocortex. Eur. J. Neurosci. 48, 2482–2497 (2018).

30. Leszczyński, M. et al. Dissociation of broadband high-frequency activity and neuronal firing in the neocortex. Sci. Adv. 6, eabb0977 (2020).

31. Nir, Y. et al. Coupling between Neuronal Firing Rate, Gamma LFP, and BOLD fMRI Is Related to Interneuronal Correlations. Curr. Biol. 17, 1275–1285 (2007).

32. Logothetis, N. K., Pauls, J., Augath, M., Trinath, T. & Oeltermann, A. Neurophysiological investigation of the basis of the fMRI signal. Nature 412, 150–157 (2001).

33. Conner, C. R., Ellmore, T. M., Pieters, T. A., DiSano, M. A. & Tandon, N. Variability of the Relationship between Electrophysiology and BOLD-fMRI across Cortical Regions in Humans. J. Neurosci. 31, 12855–12865 (2011).

34. Johnson, E. L. et al. A rapid theta network mechanism for flexible information encoding. Nat. Commun. 14, 2872 (2023).

35. Canolty, R. T. et al. High Gamma Power Is Phase-Locked to Theta Oscillations in Human Neocortex. Science 313, 1626–1628 (2006).

36. Alekseichuk, I., Turi, Z., Amador de Lara, G., Antal, A. & Paulus, W. Spatial Working Memory in Humans Depends on Theta and High Gamma Synchronization in the Prefrontal Cortex. Curr. Biol. 26, 1513–1521 (2016).

37. Reddy, L. et al. Theta-phase dependent neuronal coding during sequence learning in human single neurons. Nat. Commun. 12, (2021).

38. Mehta, M. R., Lee, A. K. & Wilson, M. A. Role of experience and oscillations in transforming a rate code into a temporal code. Nature 417, 741–746 (2002).

39. O’Keefe, J. & Recce, M. L. Phase relationship between hippocampal place units and the EEG theta rhythm. Hippocampus 3, 317–330 (1993).

40. Siegel, M., Warden, M. R. & Miller, E. K. Phase-dependent neuronal coding of objects in short-term memory. Proc. Natl. Acad. Sci. 106, 21341–21346 (2009).

41. Hopfield, J. J. Pattern recognition computation using action potential timing for stimulus representation. Nature 376, 33–36 (1995).

42. Varela, F., Lachaux, J.-P., Rodriguez, E. & Martinerie, J. The brainweb: Phase synchronization and large-scale integration. Nat. Rev. Neurosci. 2, 229–239 (2001).

43. Fries, P. A mechanism for cognitive dynamics: neuronal communication through neuronal coherence. Trends Cogn. Sci. 9, 474–480 (2005).

44. Buzsáki, G. Rhythms of the Brain. (Oxford University Press, 2006).

45. Su, M. et al. Theta Oscillations Support Prefrontal-hippocampal Interactions in Sequential Working Memory. Neurosci. Bull. 40, 147–156 (2024).

46. Shi, L. et al. Distributed theta networks support the control of working memory: Evidence from scalp and intracranial EEG. Preprint at 10.1101/2025.08.14.670214 (2025).

47. Johnson, E. L. et al. Dissociable oscillatory theta signatures of memory formation in the developing brain. Curr. Biol. 32, 1457–1469.e4 (2022).

48. Kota, S., Rugg, M. D. & Lega, B. C. Hippocampal Theta Oscillations Support Successful Associative Memory Formation. J. Neurosci. 40, 9507–9518 (2020).

49. Goyal, A. et al. Functionally distinct high and low theta oscillations in the human hippocampus. Nat. Commun. 11, 2469 (2020).

50. Vivekananda, U. et al. Theta power and theta-gamma coupling support long-term spatial memory retrieval. Hippocampus 31, 213–220 (2021).

51. Choi, K. et al. Longitudinal Differences in Human Hippocampal Connectivity During Episodic Memory Processing. Cereb. Cortex Commun. 1, tgaa010 (2020).

52. Lega, B. C., Jacobs, J. & Kahana, M. Human hippocampal theta oscillations and the formation of episodic memories. Hippocampus 22, 748–761 (2012).

53. Aktürk, T., De Graaf, T. A., Güntekin, B., Hanoğlu, L. & Sack, A. T. Enhancing memory capacity by experimentally slowing theta frequency oscillations using combined EEG-tACS. Sci. Rep. 12, 14199 (2022).

54. Bender, M., Romei, V. & Sauseng, P. Slow Theta tACS of the Right Parietal Cortex Enhances Contralateral Visual Working Memory Capacity. Brain Topogr. 32, 477–481 (2019).

55. Wang, Y., De Weerd, P., Sack, A. T. & Van De Ven, V. Distinct effects of slow and fast theta tACS in enhancing temporal memory. Imaging Neurosci. 2, imag–2–00332 (2024).

56. König, P., Engel, A. K. & Singer, W. Relation between oscillatory activity and long-range synchronization in cat visual cortex. Proc. Natl. Acad. Sci. 92, 290–294 (1995).

57. Yarbrough, J. B., Shi, L., Chattopadhyay, K., Knight, R. T. & Johnson, E. L. One of these things is not like the others: Theta, beta, & ERP dynamics of mismatch detection. Preprint at 10.1101/2025.07.11.664390 (2025).

58. Cross, Z. R. et al. The development of aperiodic neural activity in the human brain. Nat. Hum. Behav. https://doi.org/10.1038/s41562-025-02270-x (2025) doi:10.1038/s41562-025-02270-x.

59. Johnson, E. L. et al. Dynamic frontotemporal systems process space and time in working memory. PLOS Biol. 16, e2004274 (2018).

60. Flinker, A. et al. Redefining the role of Broca’s area in speech. Proc. Natl. Acad. Sci. 112, 2871–2875 (2015).

61. Johnson, E. L. et al. Bidirectional Frontoparietal Oscillatory Systems Support Working Memory. Curr. Biol. 27, 1829–1835.e4 (2017).

62. Wen, H. & Liu, Z. Separating Fractal and Oscillatory Components in the Power Spectrum of Neurophysiological Signal. Brain Topogr. 29, 13–26 (2016).

63. Watrous, A. J., Tandon, N., Conner, C. R., Pieters, T. & Ekstrom, A. D. Frequency-specific network connectivity increases underlie accurate spatiotemporal memory retrieval. Nat. Neurosci. 16, 349–356 (2013).

64. Lachaux, J.-P., Rodriguez, E., Martinerie, J. & Varela, F. J. Measuring phase synchrony in brain signals. Hum. Brain Mapp. 8, 194–208 (1999).

65. Rissman, J., Gazzaley, A. & D’Esposito, M. Dynamic Adjustments in Prefrontal, Hippocampal, and Inferior Temporal Interactions with Increasing Visual Working Memory Load. Cereb. Cortex 18, 1618–1629 (2008).

66. Johnson, E. L., Kam, J. W. Y., Tzovara, A. & Knight, R. T. Insights into human cognition from intracranial EEG: A review of audition, memory, internal cognition, and causality. J. Neural Eng. 17, 051001 (2020).

67. Johnson, E. L., Kam, J. W. Y., Tzovara, A. & Knight, R. T. Insights into human cognition from intracranial EEG: A review of audition, memory, internal cognition, and causality. J. Neural Eng. 17, 051001 (2020).

68. Snodgrass, J. G. & Corwin, J. Pragmatics of measuring recognition memory: Applications to dementia and amnesia. J. Exp. Psychol. Gen. 117, 34–50 (1988).

69. Stolk, A. et al. Integrated analysis of anatomical and electrophysiological human intracranial data. Nat. Protoc. 13, 1699–1723 (2018).

70. Asano, E., Juhász, C., Shah, A., Sood, S. & Chugani, H. T. Role of subdural electrocorticography in prediction of long-term seizure outcome in epilepsy surgery. Brain J. Neurol. 132, 1038–1047 (2009).

71. Bastos, A. M. & Schoffelen, J.-M. A Tutorial Review of Functional Connectivity Analysis Methods and Their Interpretational Pitfalls. Front. Syst. Neurosci. 9, (2016).

72. Shirhatti, V., Borthakur, A. & Ray, S. Effect of Reference Scheme on Power and Phase of the Local Field Potential. Neural Comput. 28, 882–913 (2016).

73. Rossini, L. et al. Seizure activity per se does not induce tissue damage markers in human neocortical focal epilepsy. Ann. Neurol. 82, 331–341 (2017).

74. Oostenveld, R., Fries, P., Maris, E. & Schoffelen, J.-M. FieldTrip: Open Source Software for Advanced Analysis of MEG, EEG, and Invasive Electrophysiological Data. Comput. Intell. Neurosci. 2011, 1–9 (2011).

75. Cohen, M. X. Analyzing Neural Time Series Data: Theory and Practice. (The MIT Press, 2014). doi:10.7551/mitpress/9609.001.0001.

76. Drongelen, W. van. Signal Processing for Neuroscientists. (Academic Press, London, 2018).

77. Aru, J. et al. Untangling cross-frequency coupling in neuroscience. Curr. Opin. Neurobiol. 31, 51–61 (2015).

78. Monchy, N. et al. Functional Connectivity Is Dominated by Aperiodic, Rather Than Oscillatory, Coupling. J. Neurosci. 45, e1041252025 (2025).

79. Rutishauser, U., Schuman, E. M. & Mamelak, A. N. Online detection and sorting of extracellularly recorded action potentials in human medial temporal lobe recordings, in vivo. J. Neurosci. Methods 154, 204–224 (2006).

80. Quiroga, R. Q., Nadasdy, Z. & Ben-Shaul, Y. Unsupervised Spike Detection and Sorting with Wavelets and Superparamagnetic Clustering. Neural Comput. 16, 1661–1687 (2004).

81. Ester, M., Kriegel, H.-P., Sander, J. & Xu, X. A density-based algorithm for discovering clusters in large spatial databases with noise. in vol. 96 226–231 (1996).

82. Shukla, P. & Awasthi, P. K. An introduction to signal compression by using ‘Haar Wavelets’. J. Math. Probl. Equ. Stat. 1, 64–66 (2020).

83. Soleymankhani, A. & Shalchyan, V. A New Spike Sorting Algorithm Based on Continuous Wavelet Transform and Investigating Its Effect on Improving Neural Decoding Accuracy. Neuroscience 468, 139–148 (2021).

84. Dede, A. J. O. et al. Intra- and inter-regional dynamics in cortical-striatal-tegmental networks. J. Neurophysiol. 128, 1–18 (2022).

85. Berens, P. CircStat : A MATLAB Toolbox for Circular Statistics. J. Stat. Softw. 31, (2009).

86. R Core Team.

87. R Package ‘Circular’: Circular Statistics. Agostinelli, C., & Lund, U. (2017).

