## Supplemental Material 1-3 for "Opposing effects of slow and fast theta synchrony on working memory in the human hippocampal-orbitofrontal network"

| subID | age | sex | AR | RT | SU | HC | OFC | AMY | HC micro | AMY micro | OFC micro |
|---|---|---|---|---|---|---|---|---|---|---|---|
| S1 | 15 | M | 1.0 | 1826.79 | 0 | 3 | 0 | 0 | NA | NA | NA |
| S2 | 51 | F | 0.9 | 2518.79 | 0 | 2 | 0 | 0 | NA | NA | NA |
| S3 | 38 | M | 0.8 | 3280.55 | 0 | 0 | 4 | 0 | NA | NA | NA |
| S4 | 28 | F | 0.97 | 2233.43 | 0 | 4 | 0 | 0 | NA | NA | NA |
| S5 | 43 | M | 0.83 | 1958.78 | 0 | NA | NA | NA | NA | NA | NA |
| S6 | 38 | F | 0.93 | 2248.61 | 0 | NA | NA | NA | NA | NA | NA |
| S7 | 30 | M | 1.0 | 1619.73 | 1 | NA | NA | NA | NA | NA | NA |
| S8 | 27 | M | 0.97 | 1605.45 | 0 | 1 | 3 | 10 | NA | NA | NA |
| S9 | 26 | M | 0.84 | 6225.9 | 1 | 0 | 1 | 0 | 1 | 0 | 2 |
| S10 | 25 | F | 1.0 | 2161.08 | 0 | 9 | 5 | 8 | NA | NA | NA |
| S11 | 29 | M | 0.91 | 1934.91 | 1 | 4 | 0 | 2 | NA | NA | NA |
| S12 | 21 | F | 0.98 | 2140.69 | 1 | 1 | 5 | 6 | 2 | 0 | 1 |
| S13 | 41 | M | 0.91 | 3311.36 | 1 | 3 | 3 | 6 | 1 | 4 | 2 |
| S14 | 32 | M | 0.92 | 2366.25 | 1 | 2 | 0 | 1 | 3 | 0 | 0 |
| S15 | 54 | M | 0.92 | 1898.87 | 1 | 0 | 2 | 1 | 2 | 2 | 0 |
| S16 | 23 | M | 0.95 | 2407.95 | 1 | 1 | 6 | 2 | 0 | 1 | 0 |
| S17 | 32 | M | 0.91 | 2403.07 | 1 | 1 | 0 | 0 | NA | NA | NA |
| S18 | 32 | F | 0.73 | 1797.64 | 1 | 2 | 0 | 2 | 1 | 0 | 0 |
| S19 | 13 | F | 0.89 | 2116.11 | 0 | 4 | 0 | 0 | NA | NA | NA |
| S20 | 15 | M | 0.93 | 1738.43 | 0 | 6 | 0 | 0 | NA | NA | NA |
| S21 | 17 | M | 0.84 | 2263.64 | 0 | 0 | 0 | 3 | NA | NA | NA |

**S1. Demographic and channel information for all subjects.** Left to right: Subject (SubID), Age (years), Sex, AR (see Methods), average RT (ms), single unit (SU) coverage (1=yes, 0=no), macro-channel coverage per region, and micro-channel coverage per region for each subject included in analyses. NAs demonstrate subjects with no IRASA peaks detected for slow and/or fast theta on any channel within the region. AR and RT were rounded to the nearest 100$^{th}$ decimal place for presentation purposes in this table.

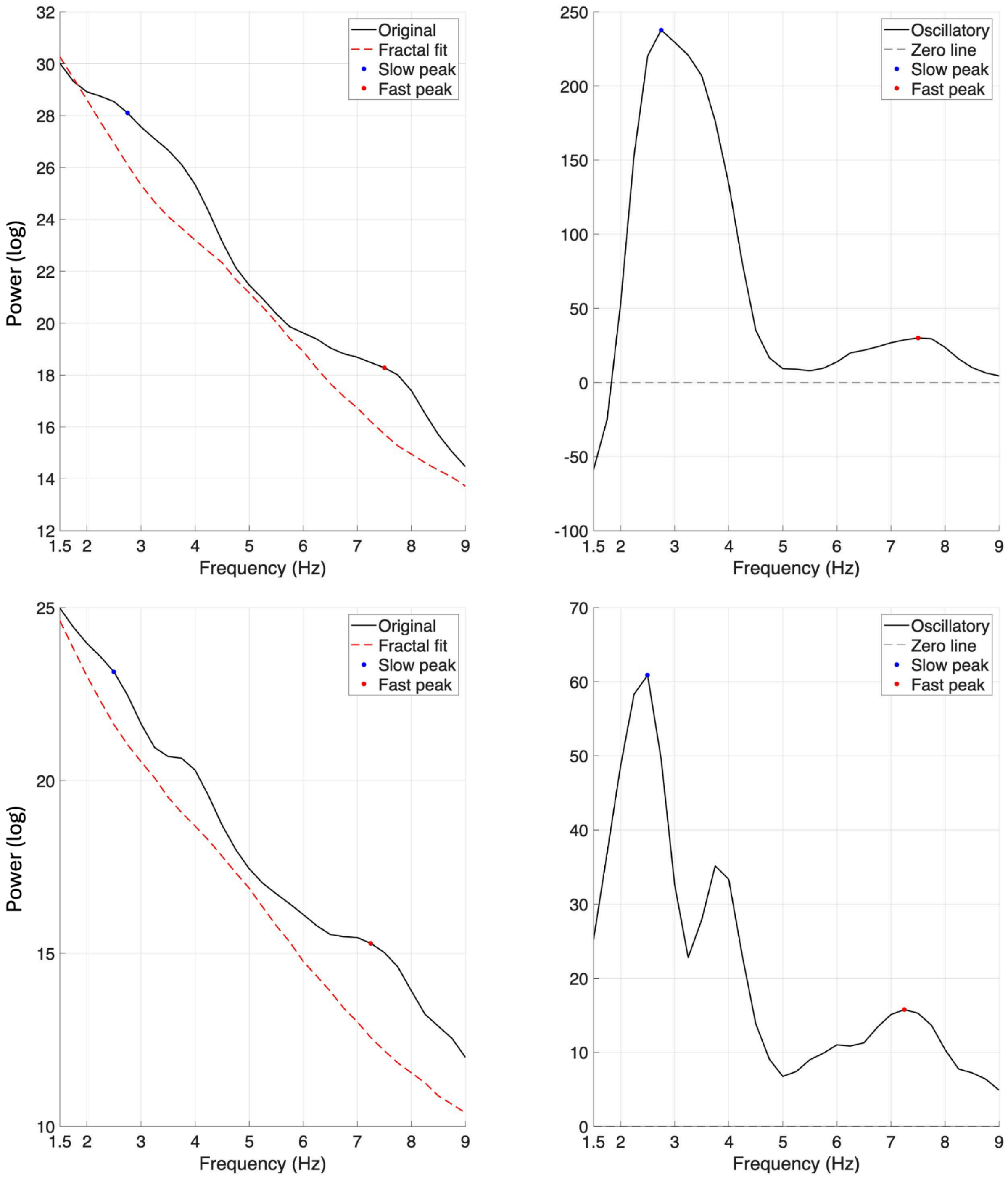


**S2. Example theta peak detection using IRASA for two representative HC channels.** For each channel (A&B), the left panel shows the power spectral density (PSD) in the theta range (1.5–9 Hz) with the aperiodic (1/f) fractal fit overlaid. The right panel shows the oscillatory component, computed by subtracting the fractal fit from the original PSD, isolating periodic activity above the aperiodic background. Detected slow theta (1.5–4.5 Hz) and fast theta (4.5–9 Hz) peaks are marked with blue and red circles respectively.

**Supplemental Text S3. Full results for preliminary single-unit phase coding analyses.**

We assessed theta phase coding of single neurons during sequence maintenance in delay 1. Theta phase coding of location, time, and concept has been previously shown in the human HC[37,39–41]. Based on this literature and our demonstration of slow and fast theta-HFB PAC in HC (see Figure 6A), we hypothesized that HC neurons would encode temporal order in the sample sequence through phase coding, wherein neurons with different stimulus preferences would fire at different theta phases. Because the speed of HC slow, but not fast, theta oscillations was modulated by task demands (see Figure 3), we anticipated that phase coding would be specific to the slow theta band. Furthermore, we hypothesized that OFC would also exhibit phase coding of single neurons to slow theta due to its parallel patterns of slow theta slowing, PAC, and PLV (see Figure 4), and the beneficial effect of slow theta HC-OFC PLV on behavioral RT (see Figure 5). We provide initial tests of this hypothesis based on the subset of 10 patients who had microwire implants in HC, OFC, and/or AMY (see Table S1). Neuronal stimulus preference was determined by the temporal position in the sample sequence that elicited the highest firing rate. Neurons that did not exhibit stimulus preference and LFPs that did not exhibit a slow or fast theta peak were excluded from analyses. After spike sorting, theta peak detection, and analysis of neuronal stimulus preference (see Methods), our sample sizes were limited and thus our results are preliminary.

We first tested our hypothesis by assessing phase coding within each group of HC neurons (i.e., respective groups of neurons exhibiting preference for stimulus 1, 2, and 3) to slow theta phase, and then repeated analyses for the other ROIs as well as fast theta. Rayleigh's test of uniformity revealed significant slow theta phase coding for HC neurons with preference for stimulus 2 ($p$=0.048, n=3) and OFC neurons with preference for stimulus 3 ($p$=0.04, n=5) (Figure S3). Slow theta phase coding was not significant for HC neurons with preference for stimulus 1 ($p$=0.96, n=4) or 3 ($p$=0.96, n=7), OFC neurons with preference for stimulus 1 ($p$=0.34, n=4), or any AMY neurons ($p \geq 0.12$). No OFC neurons exhibited stimulus 2 preference on microwires with a slow theta peak, and thus we could not test phase coding of stimulus 2. Fast theta phase coding was not significant for HC or AMY neurons ($p \geq 0.05$). No OFC no neurons exhibited stimulus preference on microwires with a fast theta peak, and thus we could not test fast theta phase coding. We then tested whether the preferred slow theta phase differed by neuronal stimulus preference within each ROI, and observed no significant effects ($p \geq 0.66$)[85].

Although these phase coding results are necessarily preliminary given sample size limitations, when taken with our demonstrations of slow theta slowing within HC and OFC (see Figure 3), synchrony between HC and OFC (see Figures 4-5), and PAC within HC and OFC (see Figure 6), our findings converge to suggest that these two regions form a slow theta circuit to encode and maintain the sequential order of the second stimuli and third stimuli in the three-stimuli sequence. We provide initial evidence that HC and OFC neurons may facilitate maintenance of sequential items via slow-theta phase coding of temporal order, agnostic to stimulus identity and spatial orientation. These findings extend previous demonstrations of HC phase coding of spatial location and conceptual information in WM[37,39,41] to HC and OFC phase coding of temporal order. These results reveal potential single neuron mechanisms for maintaining temporal order in OFC. The absence of significant fast theta phase coding, combined with our other findings, provide corroborating evidence in favor of slow theta as the primary frequency for maintaining and comparing sequential information at both local and network levels. These preliminary results require replication with larger samples, but offer initial support for slow theta phase coding as a mechanism of single neuron representations of temporal order within the HC-OFC circuit.

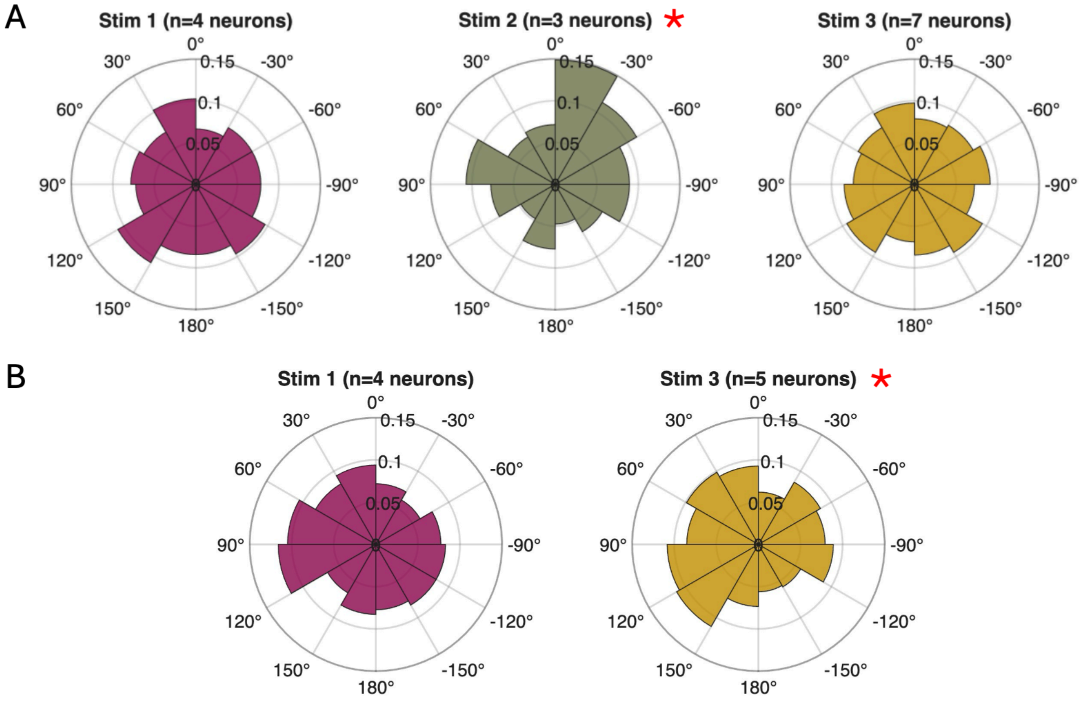


**S3. Slow theta phase locking by stimulus preference for HC (A) and OFC (B) neurons.** Polar histograms show the proportion of spikes occurring at each slow theta phase bin for neurons grouped by stimulus preference. (A) HC neurons preferring the second stimulus (green) show significant slow theta phase locking. (B) OFC neurons preferring the third stimulus (yellow) show significant slow theta phase locking.
